# The activation loop residue serine 173 of *S.pombe* Chk1 kinase is critical for the response to DNA replication stress

**DOI:** 10.1101/164244

**Authors:** Naomi Coulton, Thomas Caspari

**Author notes:** Corresponding Author: Dr Thomas Caspari, Bangor University,School of Medical Sciences, Bangor LL57 2UW, United Kingdom. Classification: BIOLOGICAL SCIENCES/Cell Biology.

## Abstract

Why the DNA damage checkpoint kinase Chk1 protects the genome of lower and higher eukaryotic cells differentially is still unclear. Mammalian Chk1 regulates replication origins, safeguards DNA replication forks and promotes fork progression. Conversely, yeast Chk1 acts only in G1 and G2. We report here that the mutation of serine 173 (S173A) in the activation loop of fission yeast Chk1 abolishes the G1-M and S-M checkpoints without affecting the G2-M arrest. Although Chk1-S173A is fully phosphorylated at serine 345 by the DNA damage sensor Rad3 (ATR) when DNA replication forks break, cells fail to stop the cell cycle. Mutant cells are uniquely sensitive to the DNA alkylation agent methyl- methanesulfate (MMS). This MMS sensitivity is genetically linked with the lagging strand DNA polymerase delta. Chk1-S173A is also unable to block mitosis when the G1 transcription factor Cdc10 is impaired. Serine 173 is equivalent to lysine 166 in human Chk1, an amino acid important for substrate specificity. We conclude that the removal of serine 173 impairs the phosphorylation of a Chk1 target that is important to protect cells from DNA replication stress.

**Summary statement:** Mutation of serine-173 in the activation loop of Chk1 kinase may promote cancer as it abolishes the response to genetic alterations that arise while chromosomes are being copied.

## Introduction

*S.pombe* Chk1 is phosphorylated in its C-terminal domain at serine-345 by Rad3 (ATR) after the kinase was recruited to a broken chromosome by Rad4 (TopBP1), Crb2 (53BP1) and the Rad9-Rad1-Hus1 ring (Capasso et al., 2002) (Lopez-Girona et al., 2001) (Saka et al., 1997) (Furuya et al., 2004). Activated Chk1 delays cell cycle progression at the G2-M boundary by stimulating Wee1 to phosphorylate the inhibitory tyrosine-15 residue of Cdc2 (CDK1) kinase and by simultaneously removing the activating tyrosine phosphatase Cdc25 from the nucleus (Furnari et al., 1999) (O’ Connell et al., 1997). It is generally believed that the Chk1 response in yeast is limited to G2 even though DNA replication forks collapse in S phase (Lindsay et al., 1998) (Francesconi et al., 1993) (Redon et al., 2003). *S.pombe* Chk1 performs a second, more enigmatic role in G1 where it prevents premature mitosis when the transcription factor Cdc10 is impaired (Carr et al., 1995). It also phosphorylates Cdc10 in the presence of methyl-methanesulfonate that alkylates the DNA template to delay G1-S transition (Ivanova et al., 2013).

Unlike in yeast, human Chk1 acts mainly during S phase. It is also phosphorylated at S345 by ATR when the kinase associates with stalled DNA replication forks via Claspin (Mrc1), in a process aided by TopBP1 (Rad4) and the 9-1-1 ring. Modification of S345 depends on the additional phosphorylation of S317 and is followed by the auto-phosphorylation of Chk1 at S296 (reviewed in (González Besteiro and Gottifredi, 2015)). This auto- phosphorylation event is important for the association of Chk1 with Cdc25A and the subsequent degradation of the phosphatase (Kasahara et al., 2010). Modification of S280 by p90 RSK kinase ensures the nuclear localisation of Chk1 (Li et al., 2012). Human Chk1 associates also with DNA lesions independently of Claspin by binding to poly-ADP-ribosyl modified PARP (Min et al., 2013). Activated Chk1 blocks late replication origins by disrupting the TopBP1-Treslin complex, promotes translesion DNA polymerases, mediate homologous recombination at broken forks through Rad51 and BRCA2, regulates fork elongation and arrests cell cycle progression by promoting the degradation of Cdc25A (reviewed in (González Besteiro and Gottifredi, 2015)). While yeast Chk1 can be deleted (Walworth and Bernards, 1996), mammalian cells depend on the kinase for viability.Interestingly, only S345 phosphorylation is required for the essential roles of Chk1 (Wilsker et al., 2008). Inhibition of human Chk1 in unperturbed cells interfers with S phase (Petermann and Caldecott, 2006) and mitosis (Zachos and Gillespie, 2007). Cdc2 (CDK1) phosphorylates human Chk1 at S286 and S301 during normal mitosis as well as in the response to DNA damage (Shiromizu et al., 2006) (Ikegami et al., 2008) with as yet unknown functional implications.

Another open question is how the catalytic activity of Chk1 is regulated. The generally accepted model predicts an auto-inhibitory complex between the N-terminal kinase domain and the C-terminal regulatory domain (Kosoy and O’ Connell, 2008) (Palermo et al., 2008). This complex is thought to open up when S345 is phosphorylated by ATR (Rad3) at sites of DNA damage. Whether this model is correct is still unclear since only the N-terminal kinase domain of human Chk1 has been crystallised (Chen et al., 2000). The activation loop adopted an open conformation in this structure which implies that Chk1 does not depend on the modification by an upstream activator as many other kinases do. How Chk1 is silenced at the end of the DNA damage response is also not fully understood. Human Chk1 is degraded after its modification at S345 in a process that is independent of the other phosphorylation sites (Zhang et al., 2005). A similar degradation does not occur in yeast. Attenuation of Chk1 correlates with its dephosphorylation at S345 by Wip1 (PPM1D) in human cells (Lu et al., 2005) and by Dis2 in *S.pombe* (den Elzen and O’ Connell, 2004). Interestingly, Wip1 is replaced by PPA2 in undamaged cells where it dephosphorylates Chk1 at S317 and S345 (Leung-Pineda et al., 2006). Currently no information is available on the regulation of Chk1 in unperturbed yeast cells.

We report here a rare separation-of-function mutation in Chk1 kinase. Mutation of serine 173 (S173A) in the activation loop of *S.pombe* Chk1 abolishes the G1-M arrest, when cells arrest at start in *cdc10* mutant cells, and the S-M arrest in the response to broken DNA replication forks. The G2-M checkpoint responses are largely intact and the mutant kinases is fully phosphorylated by Rad3. Chk1-S173 is also specifically sensitive to the alkylation of the DNA template by methyl-methanesulfonate in a manner related to the lagging strand DNA polymerase delta. We conclude that the S173A mutation impairs the activation of a downstream target of Chk1 that is specifically involved in the response to DNA replication stress. This conclusion is in line with the requirement of the equivalent lysine 166 for substrate recognition in human Chk1 (Chen et al., 2000).

## Results

### Reduced S345 phosphorylation of Chk1-S173A in unperturbed cells

Lysine 166 occupies a central position in the activation loop of human Chk1 opposite the catalytic aspartate 130 (D155 in *S.pombe*, Fig. 1A) where it may determine substrate specificity (Chen et al., 2000). The corresponding *S.pombe* residue is serine 173 (Fig. 1A) and aspartate 189 in *S.cerevisiae*.

**Fig. 1.**
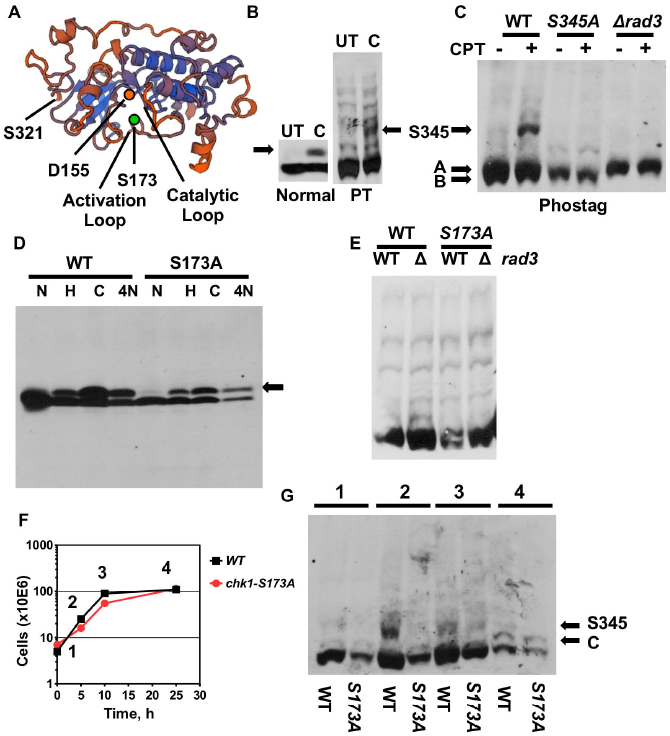
Reduced S345 phosphorylation of Chk1-S345 in unchallenged cells. (A) Model of the kinase domain of *S.pombe* Chk1. The Swiss model tool was used (swissmodel.expasy.org). The underlying crystal structure is 4czt (34.5% identity) (Chaves-Sanjuan et al., 2014). Serine-321 is the last C-terminal amino acid. (B) *chk1-HA*_*3*_ wild type cells were treated in rich medium with 10μ M camptothecin (CPT) for 3.5h at 30° C. UT = untreated. Total protein extracts were separated on normal 10% SDS page or 6% phos-tag SDS page [full image] (PT). The arrow indicates the group of shift bands related to S345 phosphorylation. (C) PT-SDS page showing extracts from *chk1-HA*_*3*_, *chk1-S345A- HA*_*3*_ and *chk1-HA*_*3*_ *rad3::ade6+* cells treated with 10μ M CPT for 3.5h [full image]. A and B indicated the hypo-phosphorylated double band. (D) Normal SDS page analysis of *chk1- HA*_*3*_ and *chk1-S173A-HA*_*3*_ cells treated with 12mM hydroxyurea (HU) and 10μ M CPT for 3.5h or with 10µM nitroquinoline 1-oxide (4NQO) for 1h at 30° C [full image] (Chk1 runs at 58kDa). (E) PT-SDS page analysing extracts from untreated *chk1-HA*_*3*_, *chk1-HA*_*3*_ *rad3::ade6+, chk1-S173A-HA*_*3*_ and *chk1-S173A-HA*_*3*_ *rad3::ade6+* cells [full image]. (F, G) Untreated *chk1-HA*_*3*_ and *chk1-S173A-HA*_*3*_ cells were grown in rich medium from a low cell number into stationary phase. Samples were withdrawn at the indicated time points and analysed on PT-SDS page [full image]. C indicates a phospho-band in stationary cells

To find out whether S173 plays a role in Chk1 activity, we mutated this residue to alanine and integrated the mutant gene with a C-terminal HA_3_ tag (*chk1-S173A-HA*_*3*_) at its endogenous locus using the Cre-*lox* recombination system (Watson et al., 2008). The integrated gene was amplified and the mutation was confirmed by DNA sequencing. We also integrated the wild type gene (*chk1-HA*_3_) (Walworth and Bernards, 1996) to exclude any effects of the flanking *lox* DNA sequences on *chk1* expression (Fig. S1).We first used the phos-tag electrophoresis assay (Caspari and Hilditch, 2015) to study the phosphorylation pattern of wild type Chk1 to establish a base line for the analysis of Chk1- S173A. Phos-tag acrylamide slows down the mobility of proteins relative to the extend of their phosphorylation (Kinoshita et al., 2006). We activated wild type Chk1 with the topoisomerase 1 inhibitor camptothecin (CPT) that breaks DNA replication forks in S phase (Pommier et al., 2010). As previously reported (Wan et al., 1999), CPT induced the mobility shift of Chk1-HA on normal SDS page which is triggered by the phosphorylation of S345 (Capasso et al., 2002) (Fig. 1B). Analysis of the same samples on a phos-tag gel revealed a larger number of phosphorylated Chk1 forms in untreated cells and a group of additional bands when cells were treated with 10μM CPT for 3.5h (Fig. 1B,C). Since these inducible bands were absent in the S345A mutant (*chk1-S345A-HA*_*3*_) (Janes et al., 2012) and in cells without Rad3 kinase (*chk1-HA*_*3*_ Δ *rad3*), they are related to the phosphorylation of serine 345 (Fig. 1C). We also noticed that the hypo-phosphorylated material of Chk1 at the bottom of the phos-tag gel consists of at least two bands (A & B in Fig 1C). Mutation of S173 to alanine (S173A) had no obvious impact on the normal band shift when cells were treated with 12mM hydroxyurea (HU), which stalls DNA replication forks, with 10μ M camptothecin (CPT) or with the UV mimetic 4-nitroquinoline 1-oxide (4-NQO) at 10μM (Fig. 1D). Also the phosphorylation pattern of Chk1-S173A in untreated cells was not significantly different from wild type (Fig. 1E).

To find out whether the unperturbed phosphorylation of Chk1 relates to cell physiology, we grew cells from early logarithmic growth into stationary phase and withdrew samples at different times (Fig. 1F). The band associated with S345 phosphorylation peaked during the most active growth phase of wild type cells (time point 2 in Fig. 1G) and was later replaced by a hypo-phosphorylated form once cells had exited the cell cycle (time point 4, band C in Fig. 1G). The peak in S345 phosphorylation reflects most likely the occurrence of endogenous DNA replication damage. It was however interesting to find that the S173A mutation lowered the amount of S345 phosphorylation during the active growth phase (time point 2 in Fig. 1G) suggesting an impaired response to replication stress.

### Chk1-S173A cells are sensitive to DNA alkylation

Since Chk1 is crucial for the G2-M checkpoint (Walworth and Bernards, 1996), we synchronised *chk1-HA*_*3*_ wild type and *chk1-S173A-HA*_*3*_ cells in G2 by lactose gradient centrifugation (Luche and Forsburg, 2009) and released them into rich medium with or without MMS (0.05%), 4NQO (10μM) or HU (12mM) at 30° C to measure the delay time. The first telling observation came when we compared the untreated strains. While wild type cells (*chk1-HA*_*3*_) entered the second cycle at around 180 min, *chk1-S173A-HA*_*3*_ cells were delayed by 20 min (Fig. 2A). Such a second cycle delay is typical for agents like CPT or HU which interfere with DNA replication (Mahyous Saeyd et al., 2014). It is therefore possible that the *chk1-S173A-HA*_*3*_ strain suffers from a DNA replication problem that triggers this short G2 delay. The UV mimetic 4-NQO and the DNA alkylation agent MMS blocked both the passage through the first G2 since DNA is instantly damaged, whereas HU caused the expected second cycle arrest as cells are only hit once they undergo DNA replication (Lindsay et al., 1998). While the S173A mutation had no impact on the HU arrest (Fig. 2B), it allowed cells to exit G2 earlier in the presence of 4-NQO and MMS (Fig. 2C, D). This partial G2-M checkpoint defect was more prominent for MMS as *chk1-S173A- HA*_*3*_ cells started to return to the cell cycle already after 80 min compared with wild type cells which arrested throughout the experiment (Fig. 2D). This checkpoint defect correlated with a high MMS sensitivity of the mutant strain (Fig. 2F, G). Interestingly, a similar loss of viability was not observed when the *chk1-S173A-HA*_*3*_ strain was treated with HU, CPT or UV light (Fig. 2E). This is an important finding as it reveals S173A as a separation-of-function mutation. MMS modifies both guanine (to 7-methylguanine) and adenine (to 3- methlyladenine) thereby inducing mismatches in the DNA that are repaired by base excision repair. Inefficient BER results in single-stranded DNA breaks independently of the cell cycle but causes DNA double-strand breaks when these gaps are encountered by a replication fork (Lundin et al., 2005). The MMS sensitivity of the *chk1-S173A-HA*_*3*_ mutant was not related to a defect in S345 phosphorylation as the mutant kinase displayed the characteristic band shift on phos-tag SDS page (Fig. 2H). Interestingly, the S345 shift was strongest at the lowest MMS concentration of 0.01% and declined at the higher concentrations.

**Fig. 2.**
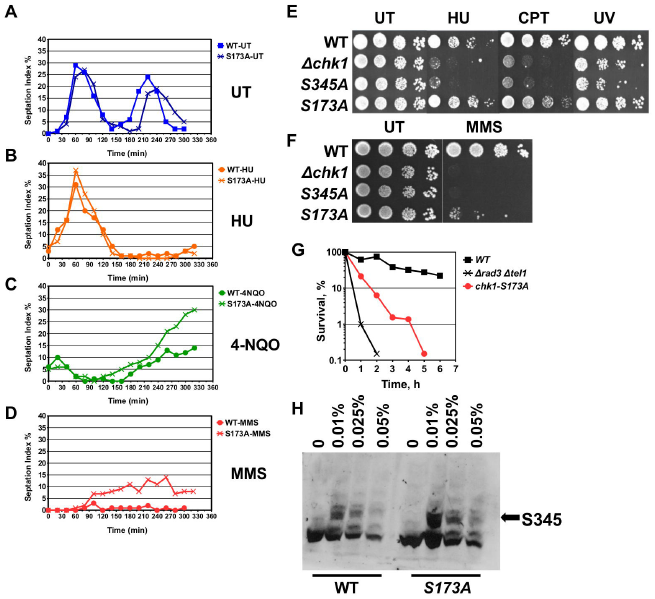
Chk1-S173A cells are MMS sensitive. (A-D). *chk1-HA*_*3*_ and *chk1-S173A-HA*_*3*_ cells were synchronised in G2 by lactose gradient centrifugation and released into rich medium containing no drug (UT), 12mM hydroxyurea (HU), 10µM nitroquinoline 1-oxide (4NQO) or 0.05% methyl-methanesulfonate (MMS). (E-F) Drop test of the indicated strains on rich medium plates containing 4mM HU, 10μM CPT, 0.01% MMS or were treated with 50J/m2 UV light (254nm). (G) Acute cell survival at 0.05% MMS. The *rad3::ade6+ tel1::leu2+* double mutant is checkpoint defective (averages of 3 repeats). (H) PT-SDS page analysis of total protein extracts from *chk1-HA*_*3*_ and *chk1-S173A-HA*_*3*_ cells treated with the indicated MMS concentrations for 3.5h at 30° C in rich medium.

### Chk1-S173A is defective in the G1-M checkpoint

In addition to its key role in G2, Chk1 blocks mitosis when *S.pombe* cells arrest at start in a *cdc10* mutant (Fig. 3A) (Carr et al., 1995). Cdc10 is a subunit of the MBF transcription factor complex that activates S phase genes during the G1-S transition (Lowndes et al., 1992). We constructed *chk1-HA*_*3*_ and *chk1-S173A-HA*_*3*_ double mutants with the temperature-sensitive *cdc10.V50* (H362Y) allele (Marks et al., 1992) and released G2- synchronised cells into rich medium at 30° C and 37° C (Fig. 3B, C). As reported previously (Carr et al., 1995), *chk1-HA*_*3*_ *cdc10.V50* cells progressed through the first cycle before arresting in G2 at the restrictive temperature of 37° C (Fig. 3B). Entry into the first cycle was delayed by 60 min due to the increase in the temperature (Janes et al., 2012). While *chk1-HA*_*3*_ *cdc10.V50* cells leaked slowly out of this G2-M arrest with only a few cells displaying the terminal cut phenotype where the new cell wall cuts through the nucleus, *chk1-S173A-HA*_*3*_ *cdc10.V50* cells entered mitosis much faster with most cells showing the cut phenoptype (Fig. 3C, D). We concuded from this experiment that the activation loop mutation impairs the G1-M checkpoint function of Chk1. Interestingly, this G1-M function of Chk1 is independent of its S345 phosphorylation as the temperature up-shift from 30° C to 37° C did not trigger the band shift on normal SDS page (Fig. 3E). Since Chk1 acts also upstream of Cdc10 to prevent entry into S phase when the DNA template is alkylated by MMS (Fig. 3F) (Ivanova et al., 2013), we synchronised *chk1-HA*_*3*_ and *chk1-S173A-HA*_*3*_ cells in metaphase using the cold sensitive *nda3.KM311* allele (Hiraoka et al., 1984) and released cells into rich medium with or without 0.01% MMS by raising the temperature from 20°C to 30°C. This experiment would allow us to measure the delay in G1-S transition induced by MMS. Untreated wild type cells (*chk1-HA*_*3*_ *nda3.KM311)* initiated DNA replication between 40 min and 60 min post-release which increased the DNA content from 2C to 4C (Fig. 3G, H). The mutant strain (*chk1-S173A- HA*_*3*_ *nda3.KM311)* showed a similar behaviour but displayed two interesting differences.Not all cells were able to escape the mitotic arrest as they maintained a 2C DNA content, and the proportion of cells that exited reached the 4C DNA content slightly earlier than wild type cells (Fig. 3H). The delayed exit from the metaphase arrest could be linked with the ability of *S.pombe* Chk1 to sustain the activation of the spindle checkpoint that delays metaphase-to-anaphase transition (Collura et al., 2005). Hence, the S173A mutation may prolong this mitotic arrest. The faster progression of the mutant cells through S phase is consistent with the reduced S345 phosphorylation during the unperturbed cell cycle (Fig. 1G) as this indicates a lower checkpoint activation. Addition of MMS delayed the accumulation of the 4C DNA content in both strains, with the S173A mutant showing a more pronounced effect (Fig. 3I). This led us to conclude that the activation loop mutation affects only the down-stream function of Chk1 that restrains mitosis in the *cdc10* mutant, but not the up-stream function which delays G1-S transition in the presence of MMS.

**Fig. 3.**
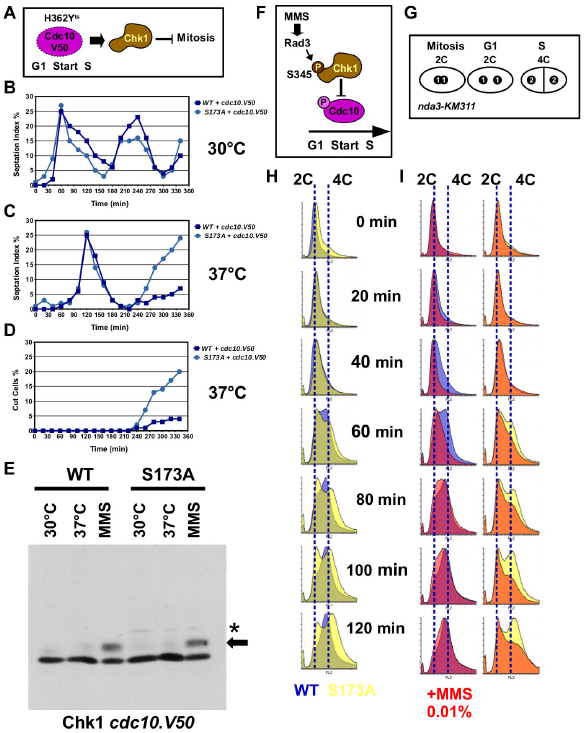
Chk1-S173A is defective in the G1-M checkpoint. (A) Inactivation of Cdc10 induces a Chk1-dependent block of mitosis. (B-D) *chk1-HA*_*3*_ *cdc10.V50* and *chk1-S173A- HA*_*3*_ *cdc10.V50* cells were grown in rich medium at 30° C, synchronised in G2 by lactose gradient centrifugation and released into rich medium at 30° C (B) or 37° C (C). Note the 60 min delay of the first cycle due to the temperature up-shift. The majority of *chk1-S173A- HA*_*3*_ *cdc10.V50* cells enter a terminal mitosis (cut phenotype) (D). (E) *chk1-HA*_*3*_ *cdc10.V50* and *chk1-S173A-HA*_*3*_ *cdc10.V50* cells were grown in rich medium at 30° C, at 37° C or at 30° C with 0.01% MMS for 3.5h. Total protein extracts were analysed on normal SDS page.(F) MMS treatment delays G1-S transition by activating Rad3-Chk1 phosphorylation of Cdc10. (G) The cold sensitive beta-tubulin gene *nda3.KM311* arrests cells with a 2x1C = 2C DNA content in metaphase. Cells reach a 4C DNA content after S phase. (H) Flow cytometry histograms of untreated *chk1-HA*_*3*_ *nda3.KM311* (blue) and *chk1-S173A-HA*_*3*_ *nda3.KM311* cells (yellow) after release from the metaphase block in rich medium. Time is post-release. (I) Flow cytometry histograms of MMS-treated (0.01%) cells after the release from the metaphase block. Dark colours = plus MMS. Blue & yellow = untreated (identical to Fig. 3H). The 2C and 4C DNA content is indicated by dotted lines.

### Chk1-S173A fails to respond to broken replication forks

The next decisive observation came when we analysed the S-M checkpoint response to broken DNA replication forks. As long as the structural integrity of a stalled fork is protected by Cds1 kinase, Chk1 activity remains low (Xu et al., 2006). Cds1 (Chk2) kinase shields stalled replication structures from nucleases and recombination enzymes (Kai et al., 2005) (Boddy et al., 2003). Chk1 is however strongly activated when forks break in the absence of Cds1, and cells without Chk1 and Cds1 are completely checkpoint defective (Lindsay et al., 1998). To test whether the S173A mutation impairs this response, we combined the *chk1-S173A-HA*_*3*_ allele with the deletion of *cds1* (Δ*cds1*). The double mutant was as HU sensitive as the Δ*chk1* Δ*cds1* strain strongly implying that the activation loop mutation blocks Chk1 activation when replication forks collapse in the absence of Cds1 (Fig. 3A). This conclusion was confirmed when we released G2-synchronised *chk1-S173A-HA*_*3*_ *Δ cds1* cells into rich medium with 12mM HU. Like the checkpoint defective *Δ chk1 Δ cds1* strain, the *chk1-S173A-HA*_*3*_ *Δ cds1* mutant entered a fatal mitosis 140min post-release (Fig. 3B). The majority of cell died while they re-entered the cell cycle indicated by the cut phenotype where one daughter cells is anuclear or where the new wall cuts through the single nucleus (Fig. 3C). Collectively, these results demonstrate an outright dependency of cells on serine 173 when replication forks break in the absence of Cds1. As in the earlier experiments, Chk1-S173A was fully phosphorylated at S345 in *Δ cds1* cells (Fig. 3D). These results imply a defect of Chk1-S173A down-stream of collapsed replication forks in the absence of Cds1.

### Chk1-S173A reduces the viability of DNA polymerase epsilon mutant cells

Because deletion of *S.pombe chk1* compromises the viability of temperature-sensitive mutants of DNA polymerase delta and epsilon (Francesconi et al., 1995), we combined mutant alleles in the three replicative DNA polymerases alpha (*swi7-H4*), delta (*cdc6.23*) and epsilon (*cdc20.M10*) with either *chk1-HA*_*3*_ or *chk1-S173A-HA*_*3*_. While testing cell growth at the semi-restictive temperature of 33° C, we noticed that the S173A mutation specifically reduced the viability of the pol epsilon (*cdc20.M10*) mutant as the *chk1-S173A- HA*_*3*_ *cdc20.M10* double mutant grew only very poorly compared to the *chk1-HA*_*3*_ *cdc20.M10* strain (Fig. 5A). DNA polymerase epsilon synthesises the leading strand (Pursell et al., 2007), is involved in long-patch BER (Wang et al., 1993), associates with the DNA replication checkpoint protein Mrc1 (Claspin) (Lou et al., 2008) and establishes heterochromatin (Li et al., 2011). The reduced viability at 33° C could suggest two roles of Chk1. Either the kinase responds to replication problems associated with the leading strand or it promotes DNA pol delta that can remove mismatches left behind by pol epsilon (Flood et al., 2015). Phos-tag analysis showed that some hypo-phosphorylated material was absent from Chk1-S173A, but this was the case for both, pol delta and epsilon (Fig.5B).

**Fig. 4.**
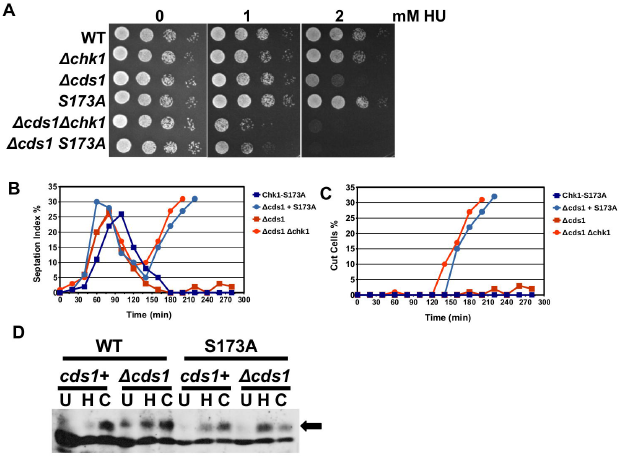
Chk1-S173A fails to respond to broken replication forks. (A) Drop test with the indicated strains at 30°C on rich medium plates. (B, C) *chk1-S173A-HA*_*3*_, *chk1-S173A- HA*_*3*_ *cds1::ura4+, cds1::ura4+* and *cds1::ura4+ chk1::kanMX4* strains were synchronised by lactose gradient centrifugation and released into rich medium with 12mM HU at 30°C (B = sepated G1/S cells; C= cut cells). (D) *chk1-HA*_*3*_, *chk1-HA*_*3*_ *cds1::ura4+, chk1-S173A- HA*_*3*_, *chk1-S173A-HA*_*3*_ *cds1::ura4+* strains were incubated for 3.5h at 30°C in rich medium [U], in 12mM HU [H] or 10μM CPT [C]. Total protein extracts were analysed on a 10% SDS page.

**Fig. 5.**
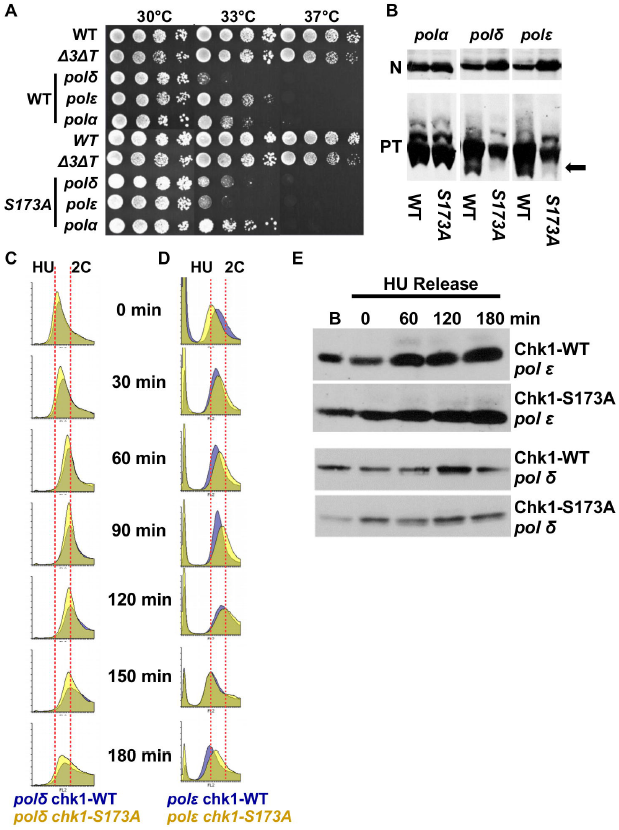
Chk1-S173A reduces the viability of DNA polymerase epsilon mutant cells. (A) Drop test with the indicated strains on rich medium plates. The *chk1-HA*_*3*_ (WT) and *chk1- S173A-HA*_*3*_ (S173A) alleles were crossed into *swi7-H4* (pol alpha), *cdc6.23* (pol delta) and *cdc20.M10* (pol epsilon). (B) PT-SDS and normal SDS (N) analysis of the strains used in the drop test. Total protein was extracted after growth in rich medium for 3.5h at 30°C. The arrow indicates the hypo-phosphorylated Chk1 protein. (C, D) *chk1-HA*_*3*_ *cdc6.23, chk1- HA*_*3*_ *cdc20.M10, chk1-S173A-HA*_*3*_ *cdc6.23* and *chk1-S173A-HA*_*3*_ *cdc20.M10* cells were synchronised in early S phase by incubating cells in rich medium with 15mM HU for 3.5h. Flow cytometry histograms were recorded at the indicated times after HU was washed out. The dotted lines indicate HU arrested and G2 (2C) cells. (E) Total protein samples were prepared from samples taken from this experiment at the indicated times and analysed on normal SDS page.

We next synchronised the strains in early S phase using the HU protocol (Luche and Forsburg, 2009) and released them back into the cell cycle to follow their progression into G2. While the S173A mutation had no impact in the case of DNA polymerase delta (*chk1- S173A-HA*_*3*_ *cdc6.23*) (Fig. 5C), it did advance cell cycle progression in the DNA polymerase epsilon strain (*chk1-S173A-HA*_*3*_ *cdc20.M10*) (Fig. 5D). The mutation in the activation loop allowed cells to acquire a G2 (2 copies, 2C) DNA content 90 min post-release, approximately 30 min earlier than the wild type Chk1 kinase (*chk1-HA*_*3*_ *cdc20.M10*). We did however find no evidence of S345 phosphorylation in any mutant strain during this experiment (Fig. 5E). The faster progression of the *chk1-S173A-HA*_*3*_ *cdc20.M10* mutant could explain why the pol epsilon strain loses viability at the semi-permissive temperature. The activation loop mutation S173A might block the phosphorylation of a down-stream target that is crucial for a reduction in leading strand synthesis when DNA polymerase epsilon is impaired or when pol delta needs to remove mismatched nucleotides.

### The MMS sensitivity of Chk1-S173A is linked with DNA polymerase delta

Given the requirement of pol delta for the removal of alkylated bases by BER (Blank et al., 1994), we tested the genetic relationship between *chk1-S173A-HA*_*3*_ and *cdc6.23*. Intriguingly, the mutation in the catalytic subunit of pol delta affected survival on MMS plates differentially depending on whether the *chk1-HA*_*3*_ wild type or *chk1-S173A-HA*_*3*_ mutant allele was present. While *cdc6.23* cells containing the wild type kinase were MMS sensitive, *cdc6.23* cells with the mutant kinase displayed some degree of resistance (Fig. 6A). We followed this observation up by conducting an acute survival test at 0.025% MMS and noticed that the *chk1-HA*_*3*_ *cdc6.23* double mutant was significantly more MMS sensitive than the pol delta (*cdc6.23*) single mutant that contains the untagged *chk1* gene (Fig. 6B). This implies that the tagged *chk1-HA*_*3*_ allele, which has been used in many studies (Walworth and Bernards, 1996), differs from the untagged gene in a *cdc6.23* mutant background. Intriguingly, the mutation in the activation loop suppressed this hyper- sensitivity to a level observed for the *chk1-S173A-HA*_*3*_ single allele (Fig. 6B). Collectively, these data show that the MMS sensitivity of the *chk1-S173A* mutation is epistatic with the *cdc6.23* mutation in the catalytic subunit of pol delta at 30° C and that the mutation also suppresses the damaging activity of the tagged wild type Chk1 kinase. The nature of this activity is as yet unknown. We suspect however that the C-terminal tag interferes with the repair function of pol delta in BER (Blank et al., 1994). To test whether the polymerase mutations interfere with S345 phosphorylation of Chk1 and Chk1-S173A, the corresponding strains were treated with 0.01% MMS at 30° C and also exposed to the semi-permissive temperature of 33° C without MMS. While both Chk1 proteins were phosphorylated at S345 in the presence of MMS, the phosphorylation of the wild type kinase was lower in the pol delta mutant coinciding with its high MMS sensitivity (Fig. 6C). Chk1 was only weakly S345 modified at 33° C in both polymerase mutants indicating that no or very little endogenous DNA damage occurs under these conditions.

**Fig. 6.**
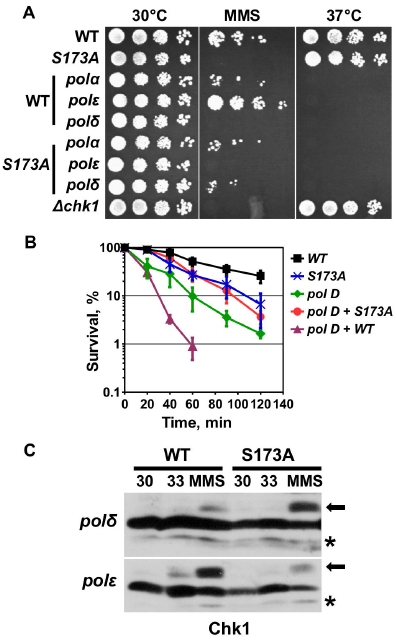
The MMS sensitivity of Chk1-S173A is linked with DNA polymerase delta. (A) Drop test with the indicated strains on rich medium plates at 30°C, 30°C with 0.01% MMS or 37°C. (B) Acute MMS survival (0.025%) at 30°C (averages and s.d. errors of 3 repeats).(C) *chk1-HA*_*3*_ *cdc6.23, chk1-HA*_*3*_ *cdc20.M10, chk1-S173A-HA*_*3*_ *cdc6.23* and *chk1-S173A- HA*_*3*_ *cdc20.M10* cells were grown at 30°C, 33°C or 30°C plus 0.01% MMS for 3.5h in rich medium. Total protein extracts were analysed on normal SDS page.

## Discussion

The only separation-of-function conditions known so far are the phosphorylation of S317 of human Chk1, which is only required for the DNA damage response but not for its essential functions (Wilsker et al., 2008), and the mutations E92D and I484T in *S.pombe* Chk1 which affect the S-M checkpoint but only at 37° C (Francesconi et al., 1997). We report here a new separation-of-function mutation, S173A in the activation loop of *S.pombe* Chk1, that abolishes the G1-M and S-M checkpoints independently of S345 phosphorylation under normal growth conditions. When *chk1-S173HA*_*3*_ cells arrest at start during the G1-S transition due to the *cdc10.V50* mutation, they cannot prevent mitosis (Fig. 3C,D). A similar problem arises when DNA replication forks break in HU medium in the absence of Cds1 (Fig. 4B, C). Since *cdc10.V50* cells arrest with unreplicated chromosomes at start (Luche and Forsburg, 2009), both Chk1 requirements must reflect distinct G1-M and S-M checkpoint activities of Chk1. What is however intriguing is that the *chk1-S173A* mutant is not CPT sensitive (Fig. 2E), although camptothecin also breaks DNA replication forks (Pommier et al., 2010). This implies that the activation loop mutation is only critical when Cds1 is absent. Since Cds1 protects damaged forks from nucleases and recombinases (Kai et al., 2005) (Boddy et al., 2003), it is possible that the activation loop mutation activates a DNA repair factor that is redundant as long as *Cds1* is active. The DNA damage signal must reach *Chk1-S173A* as the mutant kinase is phosphorylated at S345 in the presence of CPT (Fig. 1D), 4-NQO (Fig. 1D), MMS (Fig. 2H) and HU (Fig. 3D). It is therefore unlikely that the S173A mutation interferes with Rad3 activation at damaged chromosomes involving Crb2 (53BP1), Rad4 (TopBP1) and the 9-1-1 ring (Furuya et al., 2004). Since the corresponding lysine-166 in human Chk1 is involved in substrate specificity (Chen et al., 2000), it is more likely that the activation loop mutation blocks the phosphorylation of a down-stream target that is required to restrain mitosis in *cdc10* mutant cells and when forks break in *cds1* deletion cells (Fig. 7A). This target appears to be distinct from Wee1 and Cdc25 because the *chk1-S173A* strain is able to block mitosis when *cds1+* cells are treated with HU or the UV mimetic 4-NQO (Fig. 2B, C). A clear difference exists however when DNA is alkylated by MMS as *chk1-S173A* cells have a partial G2-M checkpoint defect (Fig. 2D) and are highly sensitive (Fig. 2G; 6B). The genetic link between DNA polymerase delta and Chk1-S173A may hint at this unknown target. The observation that the activation loop mutation reduces the viability of the pol epsilon (*cdc20.M10*) mutant at 33°C (Fig. 5A) could be explained by the faster progression through S phase (Fig. 5D). This faster progression may however be linked with DNA polymerase delta given that pol epsilon needs pol delta to repair any remaining mismatches in the leading strand which are not removed by its own 3`-exonuclease activity (Flood et al., 2015). Pol delta is also able to replicate the leading and the lagging strand once a fork has collapsed (Miyabe et al., 2015). The requirement of S173 for viability of the pol epsilon (*cdc20.M10*) mutant could therefore mean that Chk1 is involved in the repair activities of pol delta either when mismatched bases remain in the leading strand after MMS treatment or when the leading strand is elongated by pol delta during the homologous recombination dependent re-start of collapsed replication forks in HU-treated Δ*cds1* cells (Fig. 7A). This conclusion is strengthened by the epistatic relationship between *chk1-S173A* and *cdc6.23*, the catalytic subunit of pol delta (Fig. 6B).

**Figure 7.**
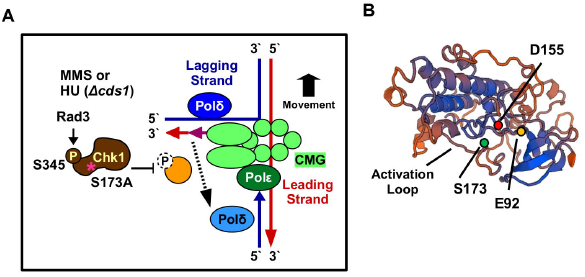
Model. (A) Chk1-S173A may be defective in the phosphorylation of a protein that allows DNA pol delta, which acts in front of the moving replication fork on the lagging strand, to remove mismatches that remain in the leading strand in the presence of MMS. Since pol delta also synthesis both strands during the recombinogenic repair of collapse forks, Chk1-S173A might also impair this function. CMG = Cdc45 + Mcm2–7 + GINS replication complex. (B) Model of the kinase domain of *S.pombe* Chk1. The underlying crystal structure is 4czt (34.5% identity).

In summary, S173A is a rare separation-of-function mutation of Chk1 that may help to dissect its role in S phase where it might link post-replication repair by DNA polymerase delta with a block over mitosis. To uncover the identity of its proposed target will however require further work. It is intriguing that one of the other known separation-of-function mutations, E92D (Francesconi et al., 1997), sits at the beginning of a loop opposite the activation loop where S173A is (Fig. 7B). The other interesting notion is that this intra-S activity of Chk1, which is not essential in yeast, may have become essential during the evolution of higher eukaryotic cells (Petermann and Caldecott, 2006).

## Materials and Methods

### Yeast strains

The genotype of the strains used in this study is *ade6-M210 leu1–32 ura4-D18*. The *rad3* gene was deleted with the *ade6+* gene and the *cds1* gene was deleted with *ura4+*. The *chk1* gene was deleted with *kanMX4* antibiotic resistance gene. *chk1-S345A-HA*_*3*_ (*h-ade6-M210 chk1::loxP-chk1-S345A-HA3-loxM leu1-32 ura4-D18); chk1-S173A-HA*_*3*_ *(h-ade6-M210 chk1::loxP-chk1-S173A-HA3-loxM leu1-32 ura4-D18)* (Fig. S1). See figure legends for further details.

### Base strain construction and integration of the Chk1 point mutations

The base strain was constructed as described in (Watson et al., 2008). The *loxP* and *loxM* Cre-recombinase recognition sequences were integrated 84nt upstream of the start codon and 84nt downstream of the stop codon (Fig. S1A) using the primers *Base-1* and *Base-2* (Fig. S1C). The point mutations S173A and S345A were introduced using fusion PCR as reported in (Janes et al., 2012). Genomic DNA from the c*hk1-HA*_*3*_ strain (Walworth and Bernards, 1996) was used as the PCR template to introduce the C-terminal HA affinity tag. The two overlapping *chk1* gene segments were amplified using the primers *Base-3* and the mutation reverse primer, and the primer *Base-4* and the mutation forward primer (Fig. S1C). The full-length fusion fragments were cloned into the *lox*-Cre integration plasmid using the restriction enzymes SphI and SacI. Integration of the mutated *chk1-HA*_*3*_ genes resulted in the loss of 4nt upstream of the start codon and of 17nt downstream of the stop codon (Fig. S1B).

### Cell synchronisation

Cells were synchronised as described in (Luche and Forsburg, 2009). HU was used at a final concentration of 15mM for 3.5h at 30 ° C in rich medium. Lactose gradients were centrifuged for 8min at 800rpm. The *nda3.KM311* mitotic arrest was performed in rich medium as reported in (Nakazawa et al., 2011). One volume of pre-warmed medium (40° C) was added to the 20° C medium to quickly raise the temperature to 30° C at the up-shift to re-start the cell cycle.

### Flow cytometry

The DNA content was measured using a CUBE 8 (Sysmex) instrument as described in (Luche and Forsburg, 2009). The histograms were produced using the free Flowing Software (http://flowingsoftware.btk.fi/)

### Phos-tag SDS page

Phostag gels (6%) were prepared and run as reported in (Caspari and Hilditch, 2015).

### Survival assays

The drop tests and acute survivals assays are described in (Kai et al., 2006).

### Antibodies

Anti-HA antibody (BioScource, Covance MMS-101P-200)

## Acknowledgements

The authours would like to thank Dr Jacqueline Hayles (The Francis Crick Institute, London, UK) for supplying the *cdc10.V50* strain.

## Author Contributions

NC performed some experiments. TC performed some experiments, designed the study and analysed the results.

